# A Population-level Strain Genotyping Method to Study Pathogen Strain Dynamics in Human Infections

**DOI:** 10.1101/2021.07.02.450905

**Authors:** Sarah J Morgan, Samantha L Durfey, Sumedha Ravishankar, Peter Jorth, Wendy Ni, Duncan Skerrett, Moira L Aitken, Edward F Mckone, Stephen J Salipante, Matthew C Radey, Pradeep K Singh

## Abstract

A hallmark of chronic bacterial infections is the long-term persistence of one or more pathogen species at the compromised site. Repeated detection of the same bacterial species can suggest that a single strain or lineage is continually present. However, infection with multiple strains of a given species, strain acquisition and loss, and changes in strain relative abundance can occur. Detecting strain-level changes and their effects on disease is challenging as most methods require labor intensive isolate-by-isolate analyses, thus, only a few cells from large infecting populations can be examined. Here we present a population-level method for enumerating and measuring the relative abundance of strains called “PopMLST”. The method exploits PCR amplification of strain-identifying polymorphic loci, next-generation sequencing to measure allelic variants, and informatic methods to determine whether variants arise from sequencing errors or low abundance strains. These features enable PopMLST to simultaneously interrogate hundreds of bacterial cells that are either cultured *en masse* from patient samples, or are present in DNA directly extracted from clinical specimens without *ex vivo* culture. This method could be used to detect epidemic or super-infecting strains, facilitate understanding of strain dynamics during chronic infections, and enable studies that link strain changes to clinical outcomes.

## Introduction

Serial culturing of chronic infection sites often repeatedly yields the same pathogen species. For instance, chronic wounds can consistently grow *Staphylococcus* and *Pseudomonas* species (1), subjects with urinary tract anomalies can be persistently infected by *Escherichia coli* (2), and chronically-infected sinuses can recurrently yield the same anaerobes (1–3). The chronic infections that afflict people with cystic fibrosis (CF) are a prime example, as the same pathogen species are frequently cultured from patients’ lung secretions for long periods. Some species, like *Pseudomonas aeruginosa* (Pa) and *Staphylococcus aureus* (Sa) can be highly abundant in the lungs of individual patients for decades or even life-long (4–9).

Repeated detection of the same bacterial species over time can imply that a single strain or lineage is continually present. However, even though most stain-level genotyping studies examine very few isolates from each infection, studies on chronic wound, urinary tract, ear, gastrointestinal, and lung infections suggest more complexity. For example, stain-level genotyping methods have shown that close to a third people with CF and Sa lung infections simultaneously harbor more than one Sa strain (4, 5, 10, 11). Likewise, up to 40% of people with CF and Pa lung infections are simultaneously infected by more two or more Pa strains (12–14), although other work has suggested lower frequencies (7, 15–19). In addition, strain relative abundance can change over time, and strains can be gained or lost in individual patients (13, 18, 20). Notorious examples are Pa epidemic strains that can infect and eventually become dominant in already-colonized patients, and markedly worsen disease (21–23).

Identifying infecting strains is important for several reasons. First, strains of the same species can differ markedly in traits like the capacity for injury, transmissibility, and resistance to antibiotics (19, 22–27). Thus, the presence of multiple strains or changes in strain relative abundance could have clinical consequences. Second, strain abundance changes could provide information about the status of host defenses, treatment efficacy, or pathogen functioning. For example, strains may recede when host defenses or treatments to which they are susceptible intensify, or when deleterious mutations arise. Likewise, new strain acquisition could indicate that host conditions have become more permissive, and analysis of succeeding strains could increase understanding of bacterial functions important *in vivo*. Third, sensitive methods for strain detect could reveal outbreaks and lapses in infection control procedures. Finally, early detection of new strains could spur eradication attempts, which may be more successful soon after strains are acquired (28–32).

Established methods for strain-level identification such as pulse-field gel electrophoresis (PFGE), multi-locus sequence typing (MLST), whole genome sequencing (WGS), and others must generally be performed on one cultured isolate at a time (16, 33–36). Because pathogen populations can be extremely large and colonies from different strains may look identical (37–41), analyzing a few colonies per sample could miss multi-strain infections and strain acquisition and loss events. Newer methods using amplification of species-specific variable regions (42, 43) are not easily adaptable to multiple pathogens, and shotgun sequencing of clinical samples (41, 44) can be limited if non-target DNA (e.g. host or other bacterial DNA) is abundant. To address these limitations, we developed PopMLST (“<u>pop</u>ulation <u>MLST</u>”), a method to enumerate and measure the relative abundance of strains present in pools of hundreds of cultured isolates, or in DNA directly extracted from clinical samples.

## Results

### Overview

In conventional MLST, bacterial colonies are isolated in pure culture, Sanger sequencing is used to identify allelic variation in MLST loci within conserved housekeeping genes (7 loci in the case of Sa and Pa), and loci allele types are determined by comparison to a database (45–47). Because a single clone is analyzed, the loci are known to be linked, in that they originate from the same bacterial isolate. Thus, loci allele identities can be combined to define the MLST type of a pure culture isolate.

In contrast, the goal of PopMLST is to enumerate the pathogen strains and measure strain relative abundance in samples that contain multiple strains and a vast excess of non-target (e.g. human) DNA. To enable this, PopMLST uses PCR to amplify MLST loci from complex samples, and next-generation sequencing to measure allele relative abundance. The PCR primers act as probes to find conserved sequences flanking MLST loci (even when the targeted species is rare), and as vectors to amplify the strain-discriminating MLST loci. Amplicons are Illumina sequenced, the bioinformatic tools are used to distinguish rare variants from errors, bin “like” sequences, and measure their relative abundance (Figure 1).

**Figure 1.**
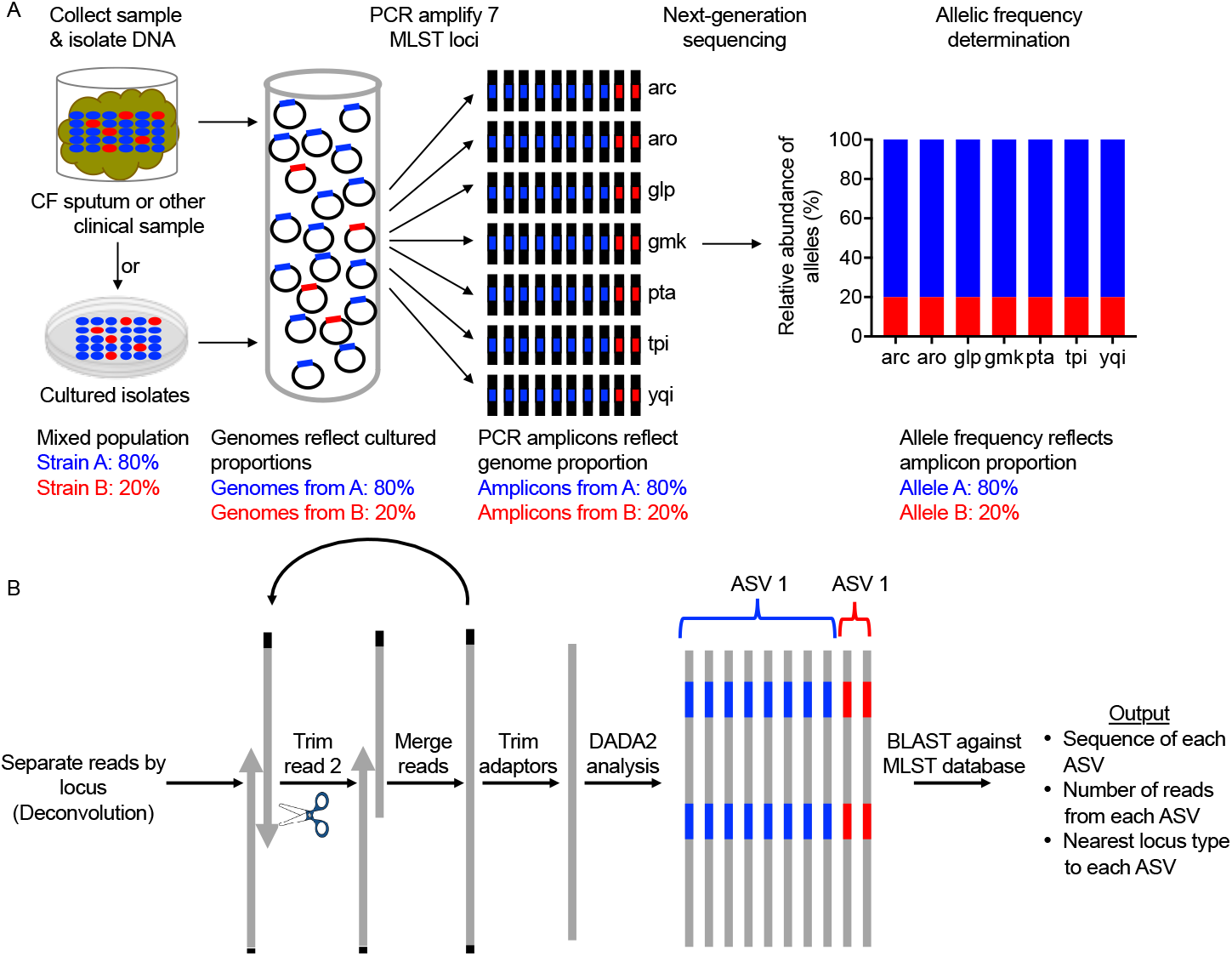
PopMLST methods. (A) PopMLST can be performed on clinical specimens (without culturing) or cultured isolates. MLST loci are PCR amplified in separate reactions, amplicons Illumina-sequenced, and the relative abundance of reads representing MLST loci are measured. (B) Bioinformatic analysis deconvolutes reads using permissive alignment to assign them to MLST loci, tests read 2 trimming lengths to optimize merging, removes adaptors, identifies amplicon sequence variants (ASVs) using DADA2, and uses BLAST to identify closest MLST locus type.

A drawback to this approach is that PCR amplification and Illumina sequencing from complex mixtures is more error-prone than Sanger sequencing of individual clones, and errors could be confused for low-abundance variant strains. We addressed this problem in several ways. (1) We used high fidelity polymerases and as few PCR cycles as possible to reduce error and PCR chimeras. (2) We adapted the DADA2 analysis pipeline (48) designed for 16S rRNA amplicon sequencing that uses statistical methods to distinguish sequencing errors from low abundance variants (Figure 1B)) (43, 48). (3) We developed bioinformatic methods to adaptively trim the lower-quality ends of the second read generated by Illumina sequencing to facilitate accurate read merging (Figure 1B and methods). (4) We amplified each MLST locus in triplicate and pooled the data to reduce random, preferential amplification of templates (i.e. “jackpot” amplifications) (49). (5) We omitted a GC-repeat rich Pa MLST loci (*aro*) that was challenging to sequence with Illumina chemistries (Figure S1 A-C) (50, 51), as we found that 91% of 3,379 MLST types in the MLST database (47) could be identified without *aro* (Figure S1D). Together these approaches mitigate, but do not fully eliminate the effects PCR and sequencing errors.

### Data interpretation

While the PCR and Illumina sequencing used in PopMLST enable analysis of complex mixtures containing multiple stains and excess non-target DNA, information from MLST loci are unlinked, as many (up to hundreds) of isolates are analyzed *en mass* and sequence reads reporting alleles from each locus are derived from separate PCR reactions. This issue does not generally limit PopMLST’s ability to enumerate and measure strain relative abundance, which can be ascertained by examining the loci with the highest number of alleles represented. This approach is effective because even though strains sometimes share MLST alleles (and PopMLST will report the sum of the shared loci’s relative abundance in these cases), the large number of alleles for each locus (e.g. Sa MLST loci have 484-892 distinct alleles, and Pa MLST loci have 137-278 distinct alleles) make it unlikely that strains would have identical alleles at enough MLST loci to prevent strain enumeration.

The MLST types of strains within mixtures can also often be determined from PopMLST data. When a limited number of strains co-exist, inference can determine which MLST alleles originate from the same strain, as linked alleles will be detected at a similar relative abundance. For example, if popMLST finds that each loci contains 3 alleles at a relative abundance of 70%: 25%: 5%, it is likely that the alleles identified at 70% relative abundance belong to one strain, alleles at 25% come from a second strain, and alleles at 5% come from a third strain. When many strains are present, if strains co-exist at similar relative abundances, or if strains happen to share several alleles, inference can fail. If knowledge of the specific MLST types is important, conventional MLST can be performed on few cultured colonies to determine which alleles are linked to one another to guide analysis of population-level data generated by PopMLST.

### PopMLST identifies single strains after *in vivo* diversification

As an initial test of the method, we performed PopMLST on pure cultures of Sa and Pa and found that >99% of reads correctly reported a single MLST type in each of 21 independent experiments (Table 1).

**Table 1.**
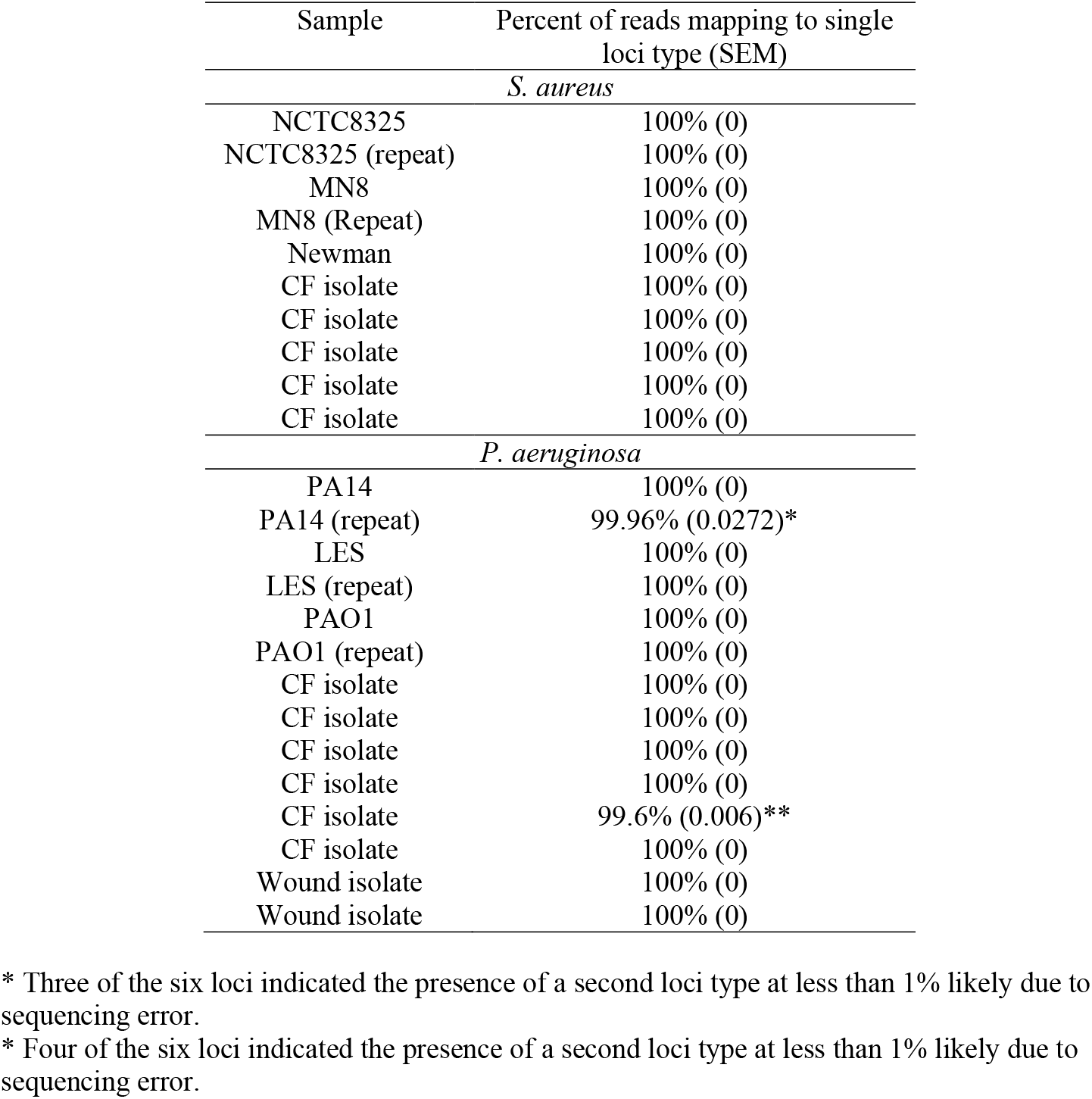
PopMLST correctly identifies single *S. aureus* and *P. aeruginosa* isolates as a single MLST type.

| Sample | Percent of reads mapping to single loci type (SEM) |
| --- | --- |
| *S. aureus* |  |
| NCTC8325 | 100% (0) |
| NCTC8325 (repeat) | 100% (0) |
| MN8 | 100% (0) |
| MN8 (Repeat) | 100% (0) |
| Newman | 100% (0) |
| CF isolate | 100% (0) |
| CF isolate | 100% (0) |
| CF isolate | 100% (0) |
| CF isolate | 100% (0) |
| CF isolate | 100% (0) |
| *P. aeruginosa* |  |
| PA14 | 100% (0) |
| PA14 (repeat) | 99.96% (0.0272)* |
| LES | 100% (0) |
| LES (repeat) | 100% (0) |
| PAO1 | 100% (0) |
| PAO1 (repeat) | 100% (0) |
| CF isolate | 100% (0) |
| CF isolate | 100% (0) |
| CF isolate | 100% (0) |
| CF isolate | 100% (0) |
| CF isolate | 99.6% (0.006)** |
| CF isolate | 100% (0) |
| Wound isolate | 100% (0) |
| Wound isolate | 100% (0) |

\* Three of the six loci indicated the presence of a second loci type at less than 1% likely due to sequencing error.
\* Four of the six loci indicated the presence of a second loci type at less than 1% likely due to sequencing error.

In CF and other chronic infections, strains genetically diversify during infection (10, 19, 25–27, 52, 53), and within-strain genetic diversity could be mistaken for strain differences. Thus, we tested PopMLST on pools of 90-96 clonally-related Pa isolates collected from different lung regions, from three CF patients undergoing lung transplantation. Whole genome sequencing showed that isolates from each subject were clonally-related to each other, but had genetically diversified via *in vivo* evolution (19). Indeed, two of the three collections exhibited hypermutator phenotypes due to mutations in either *mutL* and *mutS* mismatch repair genes (19). Core genomes of 96 isolates from the subject that was not a hypermutator contained a total 328 SNP differences, and the 96 isolates from subjects with hypermutator lineages contained 3169 and 1653 SNP differences (19).

Despite this extensive diversity, PopMLST correctly identified each of the populations as containing a single MLST type (< 0.01% of reads erroneously reported a second MLST allele) (Figure 2). These data suggest that the measures used to mitigate PCR amplification and sequencing errors are effective for pure-culture isolates and diversified clonally-related populations.

**Figure 2.**
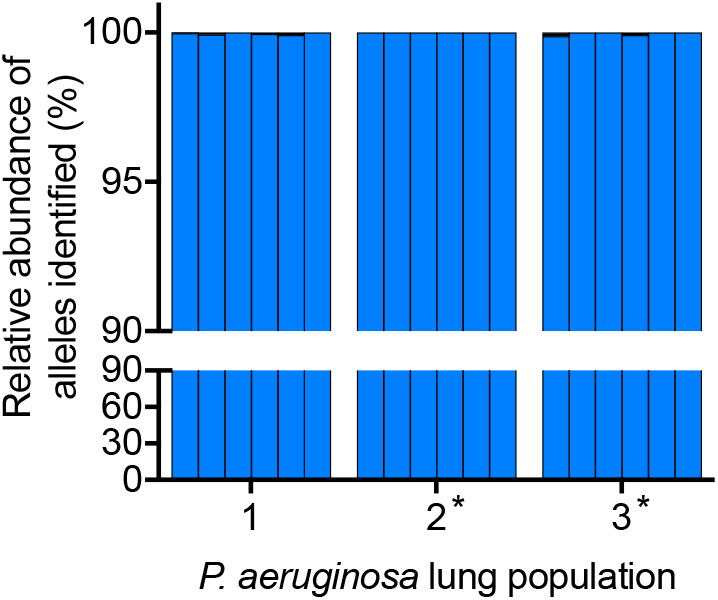
PopMLST correctly identifies genetically-diversified clonally-related Pa as a single MLST type. Plot shows the relative abundance of each MLST allele that matches the known MLST sequence (determined by genome sequencing). The six bars for each sample show the relative abundance of *acs, gua, mut, nuo, pps*, and *trp* loci (in order). Black bars indicate any additional MLST loci types detected and in all cases were less than 0.2%. * indicates hypermutable populations due to *mutS* (population 2) and *mutL* (population 3) mutations.

### PopMLST accurately measures pathogen strains in experimental mixtures

A key assumption of our approach is that the relative abundance of MLST loci present in samples is maintained through DNA extraction, amplification, sequencing, and enumeration steps (Figure 1). We therefore began testing PopMLST’s ability to detect multiple strains using defined mixtures of purified DNA from different strains. PopMLST identified the expected ratios (within 2-fold) of mixtures containing two Sa or Pa strains over a wide relative abundance range (Figure 3). Replicate experiments using different sequencing runs and different MLST types produced similar results (Figure 3A-D and S2-S3). Linear regression of data from the experimental mixtures indicated close agreement between observed and expected findings (R^2^ = 0.9916 for Sa and R^2^ = 0.9901 for Pa) with slopes approximating 1 (Sa: 1.017 [95% CI: 0.9872-1.047]; Pa: 0.9806 [95% CI: 0.9454-1.016]). PopMLST was also able to measure relative abundances of three and four strain mixtures (Figure 3E and F).

**Figure 3.**
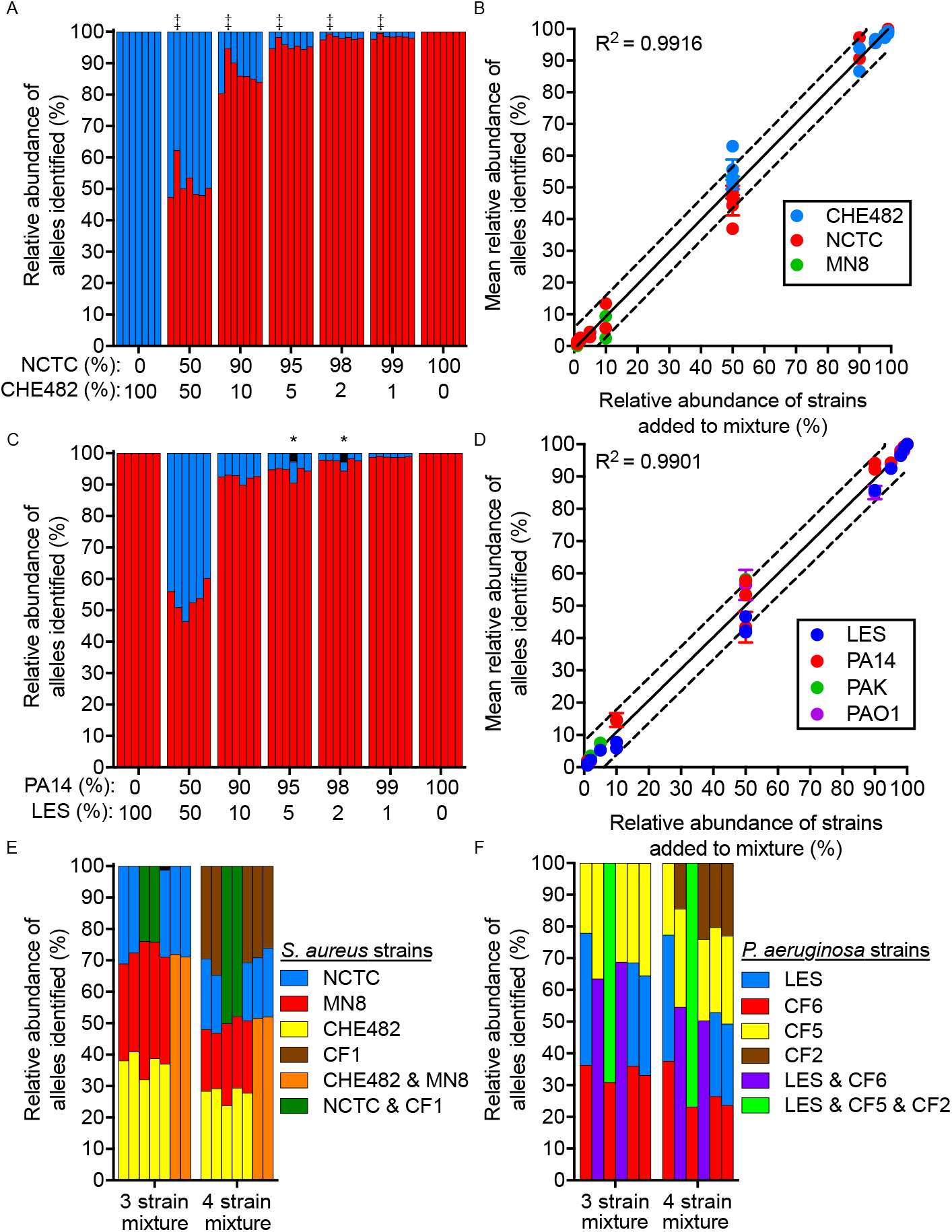
PopMLST measures strain relative abundance. (**A and C**) DNA from two Sa (**A**) and Pa (**C**) strains with different MLST types were mixed at indicated ratios. Red bars indicate reads that PopMLST called as alleles from Sa NCTC8325 (A) and Pa PA14 (C); blue bars indicate reads called as alleles from Sa CHE482 (A) and Pa LES (C). (**B and D**) shows the mean and SEM of reads mapping to indicated MLST type from 21 independent two-strain Sa mixtures (**B**) including NCTC8325 (red) with CHE482 (blue) or MN8 (green); and 20 independent two-strain Pa mixtures (**D**) of PA14 (red) and LES (blue) or PAK (green), or PAO1 (purple) and LES (blue). Data for individual alleles can be found in Figure S2 and S3. Some error bars (SEM) were smaller than symbols; solid line indicates expected result, dashed lines indicate +/- 10%. (**E and F**) Unique MLST loci alleles of the four strains tested are shown as blue, red, yellow, and brown. Alleles shared between two strains are colored as follows (red & blue = purple; yellow & red = orange; blue & brown = green (E) or blue & yellow & brown = green (F)). Bars in (A and E) show relative abundance of *arc, aro, glp, gmk, pta, tpi*, and *yqi* (in order). Bars in (C and F) show relative abundance of *acs, gua, mut, nuo, pps*, and *trp* (in order). MLST alleles identified but not present in the mixtures (likely sequencing error), are indicated in black and those detected at >1%, are indicated with *. ‡ indicates PCR bias as evidenced by one allele being consistently under or overrepresented.

Despite use of triplicate PCR reactions a single locus type was occasionally detected at higher than expected abundance (Figures 3 and S2-S3). These finding are likely due to PCR bias (indicated by ‡ in Figures 3 and S2-S3) or jackpot amplifications (indicated by # in Figures S3), which also occurs in 16S rRNA gene measurements (54) which also uses amplicon sequencing. However, because PopMLST integrates data from 6 or 7 independently-amplified loci (unlike 16S sequencing which relies on a single locus), loci that appear to be outliers can be interpreted in context of others to estimate strain relative abundance. Thus, PCR bias did not affect the number of strains detected or markedly affect strain relative abundance in the known strain mixtures (Figure 3, Figure S2-S3).

### PopMLST has a low frequency of false positive strain calls

Error inherent to PCR and Ilumina sequencing could cause PopMLST to artifactually report strains that are not present. We reexamined control experiments containing between 1-4 strains of known composition (n=38 for Pa and n=41 for Sa) to examine the effect of using different abundance thresholds to make strain presence and absence calls. As shown in Table 2, using the criterion that single variant locus be present at ≥1% relative abundance falsely registered the presence of a new strain in 9/49 (18%) of control experiments with Pa, and 7/41 (17%) of control experiments with Sa. The criterion that two or more loci be present at ≥1%, or raising the relative abundance threshold for a single allele to ≥4% produced accurate calls in all 49 Pa, and all 41 Sa experiments. We conclude that detection of a single variant loci at greater that 4% relative abundance or two variant loci at >1% relative abundance can be used as a reliable marker of strain presence.

**Table 2.** Frequency of false positive calls depending on criteria used to call MLST type presence. Accuracy is shown as function of the abundance and number of loci used to make calls.

| Abundance<br>of loci used<br>to call<br>MLST type<br>presence | *P. aeruginosa* |  | *S. aureus* |  |
| --- | --- | --- | --- | --- |
|  | 1 locus used to call<br><u>MLST type presence</u> | 2 loci used to call<br><u>MLST type presence</u> | 1 locus used to call<br><u>MLST type presence</u> | 2 loci used to call<br><u>MLST type presence</u> |
|  | # experiments with<br>false positive (%) | # experiments with<br>false positive (%) | # experiments with<br>false positive (%) | # experiments with<br>false positive (%) |
| 5% | 0/38 (0%) | 0/38 (0%) | 0/41 (0%) | 0/41 (0%) |
| 4% | 0/38 (0%) | 0/38 (0%) | 0/41 (0%) | 0/41 (0%) |
| 3% | 0/38 (0%) | 0/38 (0%) | 2/41 (5%) | 0/41 (0%) |
| 2% | 3/38 (8%) | 0/38 (0%) | 5/41 (12%) | 0/41 (0%) |
| 1% | 4/38 (11%) | 0/38 (0%) | 7/41 (17%) | 0/41 (0%) |
| 0.50% | 5/38 (13%) | 1/38 (3%) | 7/41 (17%) | 0/41 (0%) |
| 0.25% | 6/38 (16%) | 1/38 (3%) | 8/41 (20%) | 0/41 (0%) |
| 0.10% | 9/38 (23%) | 2/38 (5%) | 10/41 (24%) | 2/41 (5%) |
| 0.01% | 14/38 (37%) | 7/38 (18%) | 10/41 (24%) | 2/41 (5%) |
| 0% | 14/38 (37%) | 7/38 (18%) | 10/41 (24%) | 2/41 (5%) |

### PopMLST can detect specific MLST types with high sensitivity

In certain settings, clinicians and researchers need to detect specific strains with a known MLST type. Examples include superinfections with virulent Pa epidemic strains in people with CF already colonized by Pa, or infection control surveillance during outbreaks. Theoretically, known MLST types should be detectable with much higher sensitivity than unknown types, as it is extremely unlikely that the chance occurrence of errors would report the presence of the specific MLST loci of interest.

To test this, we measured PopMLST’s sensitivity to detect targeted low abundance MLST alleles in complex mixtures. As shown in Table 3 and Table 4, targeted low abundance alleles were detected in all experiments when present at 2% relative abundance or greater, and in almost all experiments when present at 1% and 0.1% relative abundance. These findings suggest that PopMLST could be used for early detection of known strains with high transmissibility or virulence, or to investigate efficacy of infection control measures.

**Table 3.** PopMLST’s sensitivity for detecting known *S. aureus* MLST types.

| % low<br>abundance<br>MLST type | arc | aro | glp | MLST Loci:<br>gmk | pta | tpi | yqi |
| --- | --- | --- | --- | --- | --- | --- | --- |
|  | # times loci<br>detected (range) | # times loci<br>detected (range) | # times loci<br>detected (range) | # times loci<br>detected (range) | # times loci<br>detected (range) | # times loci<br>detected (range) | # times loci<br>detected (range) |
| 10% | 4/4 (19.5-2.8) | 4/4 (6.0-1.8) | 4/4 (10.6-1.9) | 4/4 (14.0-2.8) | 4/4 (14.1-2.3) | 4/4 (14.9-2.4) | 4/4 (16.0-2.3) |
| 5% | 4/4 (5.3-3.8) | 4/4 (3.2-1.6) | 4/4 (6.1-1.8) | 4/4 (5.2-2.7) | 4/4 (4.6-3.0) | 4/4 (5.5-3.4) | 4/4 (4.7-3.9) |
| 2% | 4/4 (2.4-1.7) | 4/4 (1.2-0.6) | 4/4 (3.4-0.7) | 4/4 (11.1-2.0) | 3/4 (1.7-0) | 4/4 (2.3-1.4) | 3/4 (1.9-0) |
| 1% | 3/4 (2.2-0) | 4/4 (1.1-0.1) | 3/4 (1.5-0) | 4/4 (1.5-0.2) | 2/4 (1.4-0) | 3/4 (1.5-0) | 3/4 (2.0-0) |
| 0.1% | 2/3 (0.4-0) | 3/3 (0.3-0.1) | 2/3 (0.3-0) | 3/3 (1.5-0.2) | 0/3 (0-0) | 3/3 (0.3-0.2) | 1/3 (0.2-0) |

**Table 4.** PopMLST’s sensitivity for detecting known *P. aeruginosa* MLST types.

| % low<br>abundance<br>MLST type | MLST loci: |  |  |  |  |  |  |  |  |  |  |  |
| --- | --- | --- | --- | --- | --- | --- | --- | --- | --- | --- | --- | --- |
|  | acs |  | gua* |  | mut |  | nuo* |  | pps |  | trp |  |
|  | # times loci<br>detected (range) |  | # times loci<br>detected (range) |  | # times loci<br>detected (range) |  | # times loci<br>detected (range) |  | # times loci<br>detected (range) |  | # times loci<br>detected (range) |  |
| 10% | 4/4 | (13.5-6.2) | 3/3 | (24.8-4.1) | 4/4 | (14.9-6.9) | 3/3 | (13.3-8.2) | 4/4 | (14.8-3.9) | 4/4 | (15.2-6.0) |
| 5% | 3/3 | (7.0-4.6) | 2/2 | (6.6-4.8) | 3/3 | (7.7-2.4) | 1/2 | (6.6-1) | 3/3 | (7.6-4.6) | 3/3 | (7.7-2.8) |
| 2% | 3/3 | (3.18-1.7) | 2/2 | (2.3-2.1) | 3/3 | (3.2-1.4) | 2/2 | (2.8-1.9) | 3/3 | (4.2-1.7) | 3/3 | (3.5-2.0) |
| 1% | 4/4 | (1.6-0.7) | 3/3 | (4.0-0.4) | 4/4 | (2.1-0.7) | 3/3 | (1.6-1.1) | 4/4 | (2.7-0.5) | 4/4 | (1.8-0.7) |
| 0.1% | 3/3 | (0.3-0.1) | 2/3 | (0.5-0) | 3/3 | (0.1-0.06) | 1/3 | (0.4-0) | 3/3 | (0.3-0.08) | 3/3 | (0.2-0.08) |

\* Two of the strains used for one of the 4 mixing experiments shared alleles at these loci, because the two strains could not be differentiated at these loci, they were eliminated from analysis for these loci.

### PopMLST works in the presence of excess human or non-target bacterial DNA

Clinical samples can contain vast amounts of human and non-target bacterial DNA. For example, despite high pathogen density (Pa can reach 10^8^-10^9^ CFU/ml in CF sputum), 95-99% of CF sputum DNA is human (55), and DNA from other pathogens or oral bacteria can also be highly abundant.

We investigated the effects of contaminating DNA on PopMLST two ways. First, we preformed PCR on human and non-target bacterial DNA (including closely-related species) using Sa and Pa PopMLST primers. Amplicon yields and the number of reads mapped to Sa and Pa MLST loci in these experiments were similar to no-template controls (Figure 4A-D). Second, we tested the ability of PopMLST to detect strains in the presence of 95% human DNA, and found that the vast excess of human DNA did not compromise detection, even when strains were present at as low as 1% relative abundance (Figure 4E-F).

**Figure 4.**
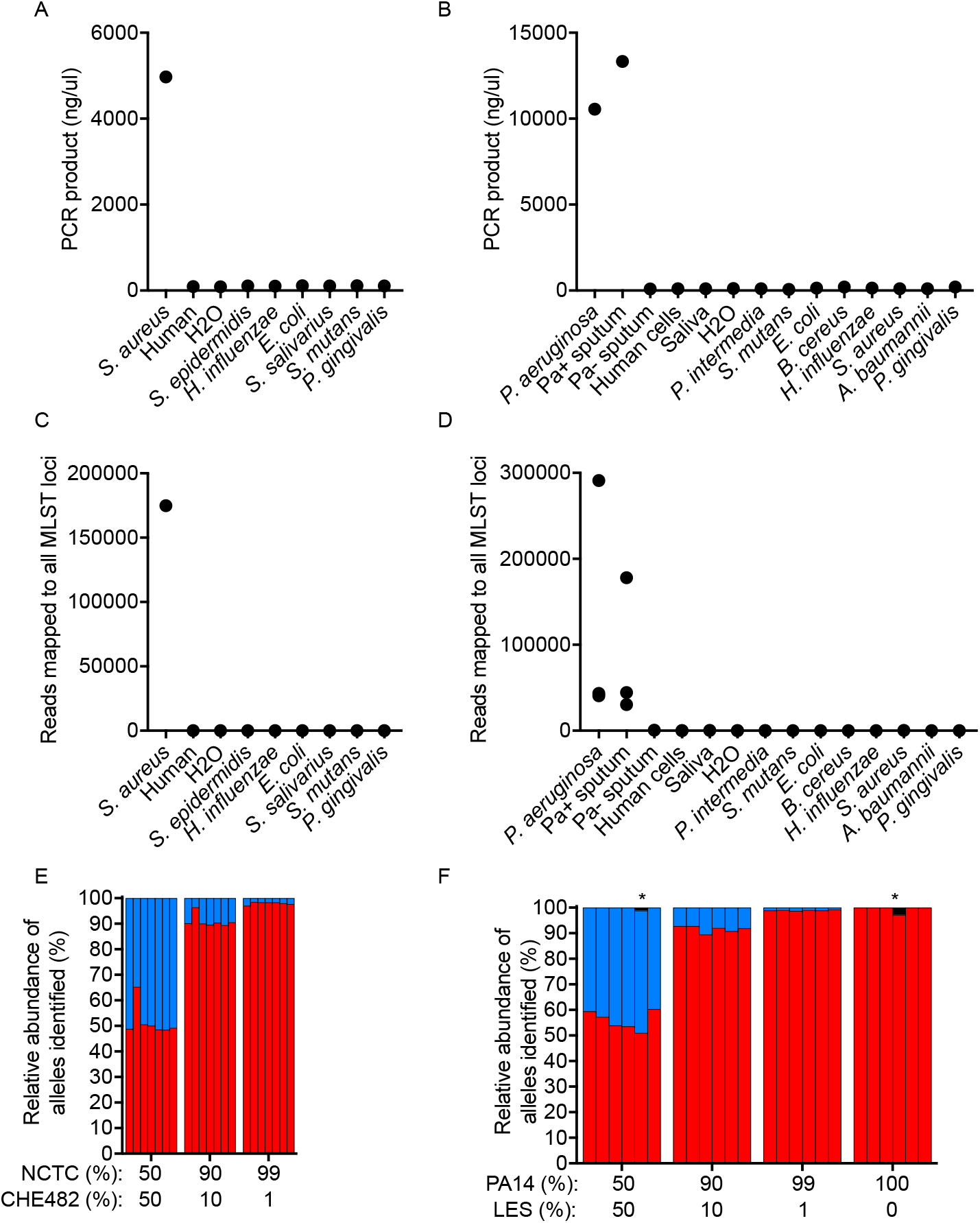
Heterologous DNA does not interfere with PopMLST. Concentrations of product after PCR using PopMLST primers for Sa (**A**) and Pa (**B**) on DNA from the indicated sources; ‘human’ indicates DNA extracted from tissue culture cells; ‘H2O’ indicates ultrapure water; ‘Pa+ sputum’ indicates sputum from a CF subject culture-positive for Pa; and ‘Pasputum’ indicates sputum from a CF subject culture-negative for Pa. The sum of reads produced by PopMLST that mapped to seven Sa (**C**) or six Pa (**D**) MLST loci are shown for samples containing target and non-target DNA from PCR reactions in A and B. (**E and F**) 95% human DNA from tissue culture cells was added to the same mixtures of two control strains from Figure 2C. (**E**) Bars for each mixture show relative abundance *arc, aro, glp, gmk, pta, tpi,* and *yqi* matching the MLST type of NCTC8325 (red) or CHE482 (blue). (**F**) Bars for each mixture show relative abundance of *acs, gua, mut, nuo, pps*, and *trp* (in order) matching the MLST type of PA14 (red) or LES (blue). * indicates the presence of an unexpected loci type (black), likely due to sequencing error.

### PopMLST measures strain abundance in clinical samples

Encouraging results with experimental strain mixtures led us to perform proof-of-principle tests of PopMLST on clinical samples. In the first test, we cultured sputum from seven Sa-infected CF subjects, and performed PopMLST on DNA prepared from ~100 colonies that grew from each sample (scraped *en mass* from culture plates). PopMLST reported that three of seven samples contained two Sa MLST types (Figure 5A). We tested these results by Sanger sequencing a distinguishing MLST locus in 20-30 individual colonies from each sample, and found MLST types at a similar relative abundance as determined by PopMLST (R^2^ = 0.9247; slope = 0.9296 [95% CI: 0.5612-1.298]) (Figure 5A-B).

**Figure 5.**
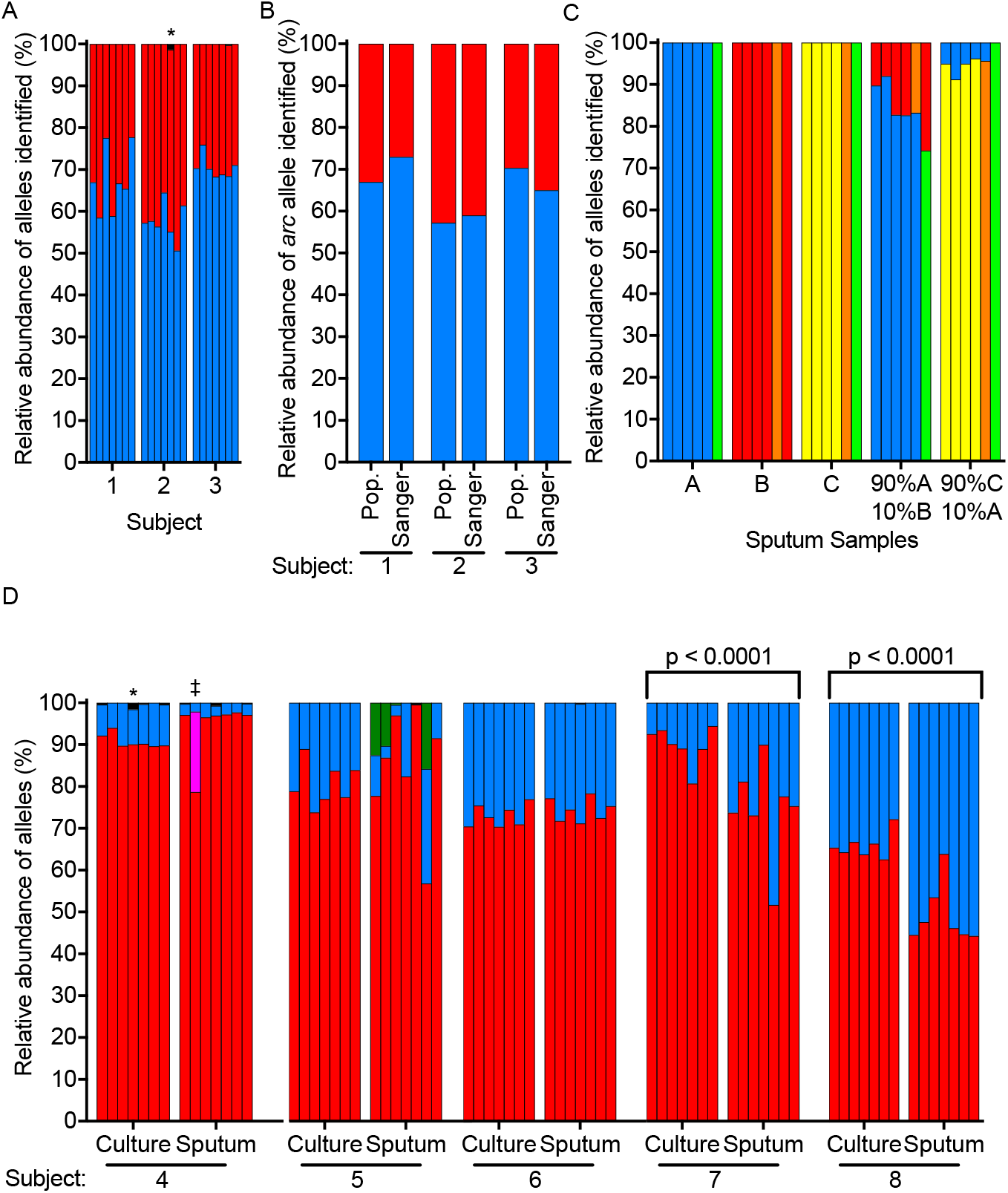
PopMLST measures MLST types in clinical samples. **A**. PopMLST was performed on ~95 Sa isolates cultured from 3 CF subjects, blue and red bars indicate different MLST loci alleles. The seven bars for each sample show relative abundance of *arc, aro, glp, gmk, pta, tpi*, and *yqi* (in order). The *pta* locus showed a third allele (black bar) which was 1 nucleotide different than the predominant allele and likely represents sequencing error. **B**. The relative abundance of the *arc* locus as measured by PopMLST (Pop) and by individually Sanger sequencing (Sanger) the *arc* locus of 20-30 isolates from each sample represented in A. **C**. PopMLST performed directly on DNA isolated from sputum (samples A, B, C) and from indicated mixtures of these samples. Red, blue, and yellow bars indicate the abundance of MLST alleles corresponding to sample A, B, and C respectively, with green bars indicating an allele shared between A and C and orange bars indicating an allele shared between B and C. Control experiments examining >100 Pa isolates cultured from sputum A, B, and C showed each contained a single Pa MLST type. **D**. Sa PopMLST of scrapes of >100 colonies (culture) and directly from sputum (sputum). Red and blue indicate allele frequencies of the predominant and secondary MLST types, respectively. Sputum of subject 5 contained three loci from an additional allele, likely indicating a third un-cultured MLST type (green). Presence of the MLST types shown in red and blue were confirmed by MLST typing of single isolates. Significant difference (p < 0.01, by multiple t-test) in abundance of the minor allele across the 7 loci is indicated by the p-value. ‡ indicates detection of a third loci allele with a single SNP different than the predominant loci, likely due to either sequencing error or mutation; * indicates the presence of a third allele (black), likely due to sequencing error.

Second, we tested PopMLST directly on CF sputum (without culturing isolates) by mixing two sputum samples that were known to harbor different Pa strains. Each sputum sample contained a single strain by popMLST, and both strains were detected in the mixtures of the two sputum samples (Figure 5C).

Finally, we analyzed sputum samples from 5 subjects that were all known to harbor 2 Sa strains each. We performed PopMLST on DNA prepared directly from sputum and from ~100 Sa isolates from each sample scraped *en mass* from culture plates, and compared results to Sanger sequencing select MLST loci from individual cultured isolates (as above). As shown in Figure 5D, PopMLST detected the same MLST types in sputum DNA and culture scrapes from all 5 subjects. In two subjects, strain relative abundance differed significantly in sputum as compared to the culture scrapes (Subject 4 and 5 p < 0.001 by multiple t-test). This finding could be due to differing sensitivity of strains to host defenses (i.e. more of one strain’s DNA originates from dead cells), or differential growth capacity of strains in *ex vivo* culture conditions.

## Discussion

Infecting bacteria can exhibit heritable diversity at the species, strain, and intra-strain level as diversity can evolve within lineages during infection. While recent findings and new methods have accelerated work on species diversity (56–64) and the diversification of a single strain (10, 19, 25–27, 52, 53), studies of strain-level diversity have lagged. A major factor is that established methods to detect strains must generally be performed on one cultured isolate at a time (16, 33–36). Thus, non-dominant strains are difficult to detect.

PopMLST addresses this limitation, as it can estimate the relative abundance of strains in pools of tens to hundreds of isolates that have been cultured from infected sites, and can be used on DNA extracted directly from clinical specimens without prior culture. Other advantages include inherent technical approaches to minimize PCR and sequencing error, its robustness when human or other bacterial DNA is in vast excess (Figure 4), its ability to detect targeted strains at very low relative abundance (Table 3 and 4) and strains with *in vivo* growth defects, when the method is used on DNA prepared directly from clinical specimens. Furthermore, the method is accurate even when intra-strain genetic diversity has evolved (Figure 2), likely because MLST loci are within conserved “housekeeping” genes that are less variable than elements involved in Spa typing (6), RAPID or PFGE (16, 33). Finally, MLST databases are in widespread use and exist for over 100 bacterial species (47), so PopMLST can be easily be adapted for use for many organisms and results are comparable between laboratories.

PopMLST also has several limitations. First, PopMLST cannot distinguish between unrelated strains having the same MLST sequence type (11, 35), although co-infection with such strains is unlikely for pathogens for which many MLST types have been identified (there are ~3,500 Pa and ~ 5,500 Sa described MLST types to date (47)). Second, while PCR and Illumina sequencing enable the method to be used on complex mixtures containing multiple strains and abundant nontarget DNA, these techniques are subject to errors and biases. We reduce, but cannot entirely eliminate, the effect of these problems using replicate PCR, adaptive trimming, and statistical methods which enable strains to be detected at ~1-3% relative abundance. Third, PopMLST does not identify which MLST loci are linked in individual isolates. While this limitation does not compromise strain enumeration and relative abundance measurements under most circumstances, it can sometimes complicate identification of strain types present.

We expect that the use of PopMLST could increase understanding of strain dynamics during infections. Patients with chronic bacterial infections frequently experience highly variable disease manifestations, treatment responses, and rates of progression. Such variation can be seen between individual patients infected with the same pathogen(s), or in individual subjects at different times even when no change in infecting pathogen species occurs. The acquisition of different strains of a given species or changes in strain relative abundance could account for some of this variation (65). PopMLST will enable tests of new hypotheses exploring the effects of strain-level diversity on treatment resistance and disease manifestations. The methods could also be useful for detecting infection outbreaks in already colonized patients, for testing the adequacy of infection control procedures.

## Methods

### Patient samples

Sputum samples were collected in accordance with University of Washington Institutional Review Board (protocols numbers 06-4469 and STUDY00011983), and St. Vincents Hospital, Dublin IE (RS20-048). Patients provided written informed consent prior to collection of samples. Sa was isolated after sputolysin diluted sputum was cultured on Mannitol Salts Agar (Difco). Populations were scraped from plates containing >100 colonies by flooding the plate with 2 ml of LB and using a L-spreader to resuspend the bacteria. Pa was isolated after sputolysin diluted sputum was cultured on MacConkey (Difco). All cultures were stored at −80°C in 15% glycerol prior to analysis.

### DNA isolation

DNA extraction of Sa samples was performed using the DNeasy Power Soil Kit (Qiagen) with the following modifications: samples were incubated with 2.9 mg lysozyme and 0.14mg lysostaphin prior to lysis with 0.1 mm beads using bead beater (Mini-Beadbeater-16; Biospec) (41). Pa DNA was isolated from 100 ul of resuspended culture using the DNeasy Blood and Tissue kit (Qiagen) using the protocol for gram negative bacteria. DNA was isolated from 100 ul of fresh sputum using the Microbiome Kit (Qiagen).

### Control mixtures

Control strains (Table S1 and Table S2) were streaked from freezer stocks and an isolate was grown overnight prior to DNA isolation. Cultured HELA or HEK293 cells were pelleted and DNA was isolated as above. Isolated DNA was quantified by Qubit, and mixed at ratios described in the figures. MLST types of control strains were based on the MLST database and confirmed by MLST typing of single isolates if necessary (Table S1 and Table S2). PAO1-lacZ:PA14 mixtures were pre-mixed at designated ratios and plated on LB+xGal to confirm the ratio. Growth from the plate was scraped and subjected to DNA isolation as above.

### Amplification and sequencing with PopMLST

5 ng/ul DNA from cultured bacteria or 20 ng/ul DNA direct from sputum or from bacteria mixed with human DNA was amplified by PCR using published MLST primers for Sa and Pa (45, 46) with Illumina adapters on the 5’ ends to enable next generation sequencing of MLST loci (Table S3). PCR amplification of each of the seven MLST loci is performed in triplicate to reduce chances of random PCR bias using reagents listed in Table S4. Triplicate reactions were pooled after PCR and amplified DNA was visualized by agarose gel electrophoresis and quantified by Pico green (Thermo Fisher). After cleaning with Ampure beads (Beckman Coulter), the seven MLST loci for each sample were pooled in equimolar amounts and barcoded with Illumina Nextera XT indexes. PCR amplification, indexing, and cleanup was performed as described in the 16S Metagenomic Sequencing Library Preparation guide (Illumina). Barcoded MLST loci were sequenced on the Illumina MiSeq to produce 2 x 300 bp paired-end reads.

### Bioinformatic analysis

Methods outlined below for PopMLST will be available at https://github.com/marade/PopMLST upon publication. Reads were deconvolved based on their locus-specific primer sequence using Python tre, with approximate matching to MLST loci allowing for up to a 25% mismatch (https://github.com/laurikari/tre/). The 3’ end of read 2 was trimmed using a binary search algorithm designed to maximize the number of merged reads of the correct size. Trimmed reads for all MLST loci, except *yqi*, were merged using VSEARCH 2.13.4 fastq_mergepairs. Two basepairs of the sequence in the *yqi* locus beyond the 3’ ends of read 1 and 2 (due to the length of this amplicon) were artificially supplied (these bases are conserved according to the MLST database (47)). *yqi* reads were joined using VSEARCH 2.13.4 fastq_join. Merged reads, with their adaptors trimmed using Cutadapt 2.3, were then processed using DADA2 to generate amplicon sequence variants (ASVs) for each locus.

To determine the identity and quantify the relative abundance of each MLST locus, the ASVs were queried against the PubMLST database (https://pubmlst.org/saureus/ and https://pubmlst.org/paeruginosa/) (47) for the appropriate species using BLAST+ BLASTN (66). The matching sequence with the highest identity and longest length (less than or equal to the maximum locus length present in the database) was used to label each ASV by locus type, with less than 100% identity matches being marked as potential novel alleles. The resulting output table includes each MLST loci type identified, the ASV, and the number of reads assigned to each type, much like a classic 16S OTU table.

## Author Contributions

Conceptualization, S.M., S.D., and P.S; methodology, S.M., S.D., and M.R.; software, M.R.; validation, S.M. and S.D.; investigation, S.M., S.D., S.R., and D.S.; resources, M.A., S.S., P.J., E.M.; writing, S.M., S.D., and P.S.; supervision, PS.; funding acquisition, P.S. All authors have read and agreed to the published version of the manuscript.

## Acknowledgments

The authors would like to thank Hillary Hayden and Michael Parkins for critical review of the manuscript. The authors would like to thank the laboratories of Matthew Parsek and Lucas Hoffman for providing strains.

