## Supplemental Figures and Tables for "A Population-level Strain Genotyping Method to Study Pathogen Strain Dynamics in Human Infections"

### Supplemental Data

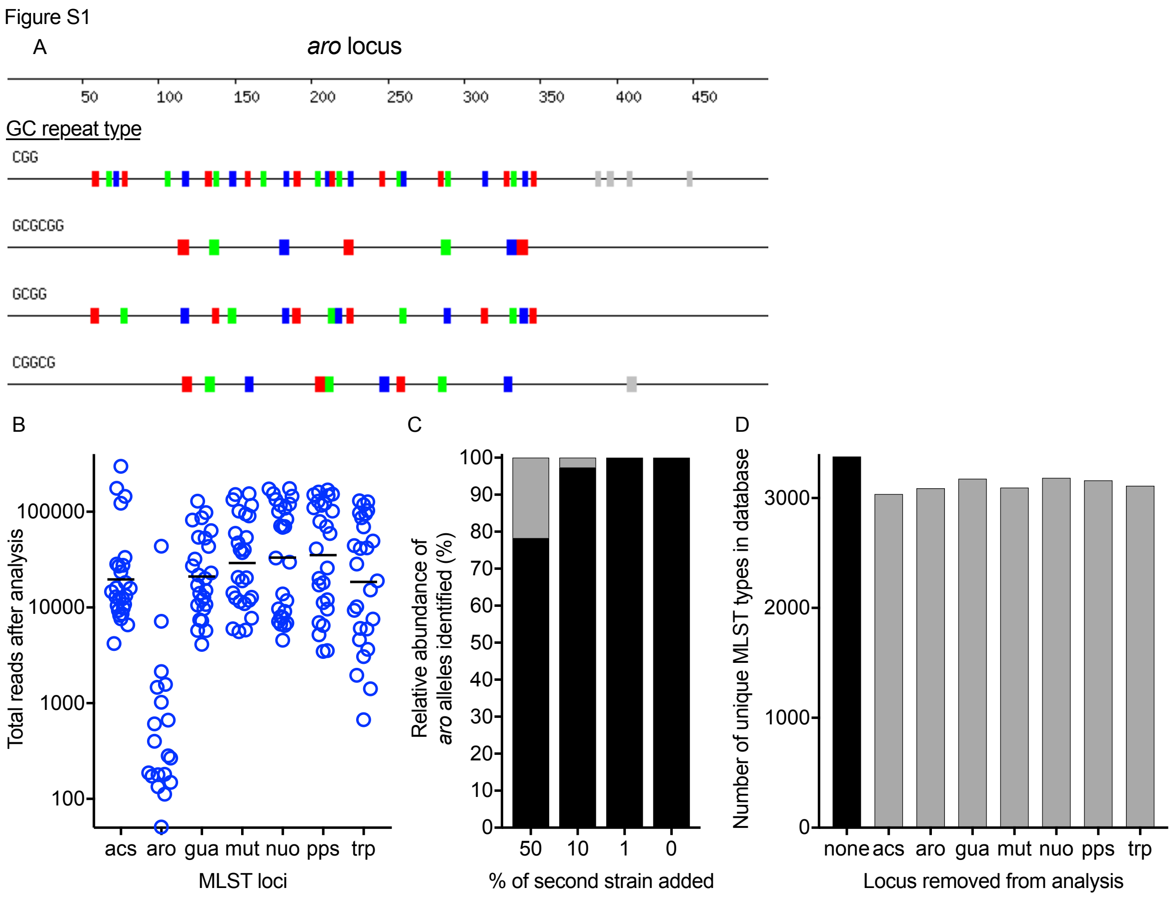


**Figure S1. *P. aeruginosa aro* MLST loci are underrepresented by PopMLST. A.** Schematic representation of high GC repeats in the *aro* MLST locus with each box showing the location of repeats identified by Geneious Prime 2020.1.1 (https://www.geneious.com). **B.** The number of reads recovered from *aro* are more than a log lower than other loci despite four-times higher levels of DNA used as input for Ilumina sequencing. **C.** Despite the low number of reads as shown in B, the *aro* reads detected were in ratios are similar to the expected ratios of the strains and the ratios of the other loci shown in Figure 3C. **D.** Analysis of the 3391 MLST types in the *P. aeruginosa* MLST database shows that omitting any one of the seven MSLT loci (indicated on the x-axis) still enables most MLST types to be identified. The number of unique combinations of MLST allele types when all seven alleles were utilized is indicated in black. The alleles for a given locus were removed and the analysis for unique combinations of allele types using the remaining six loci were determined (grey bars).

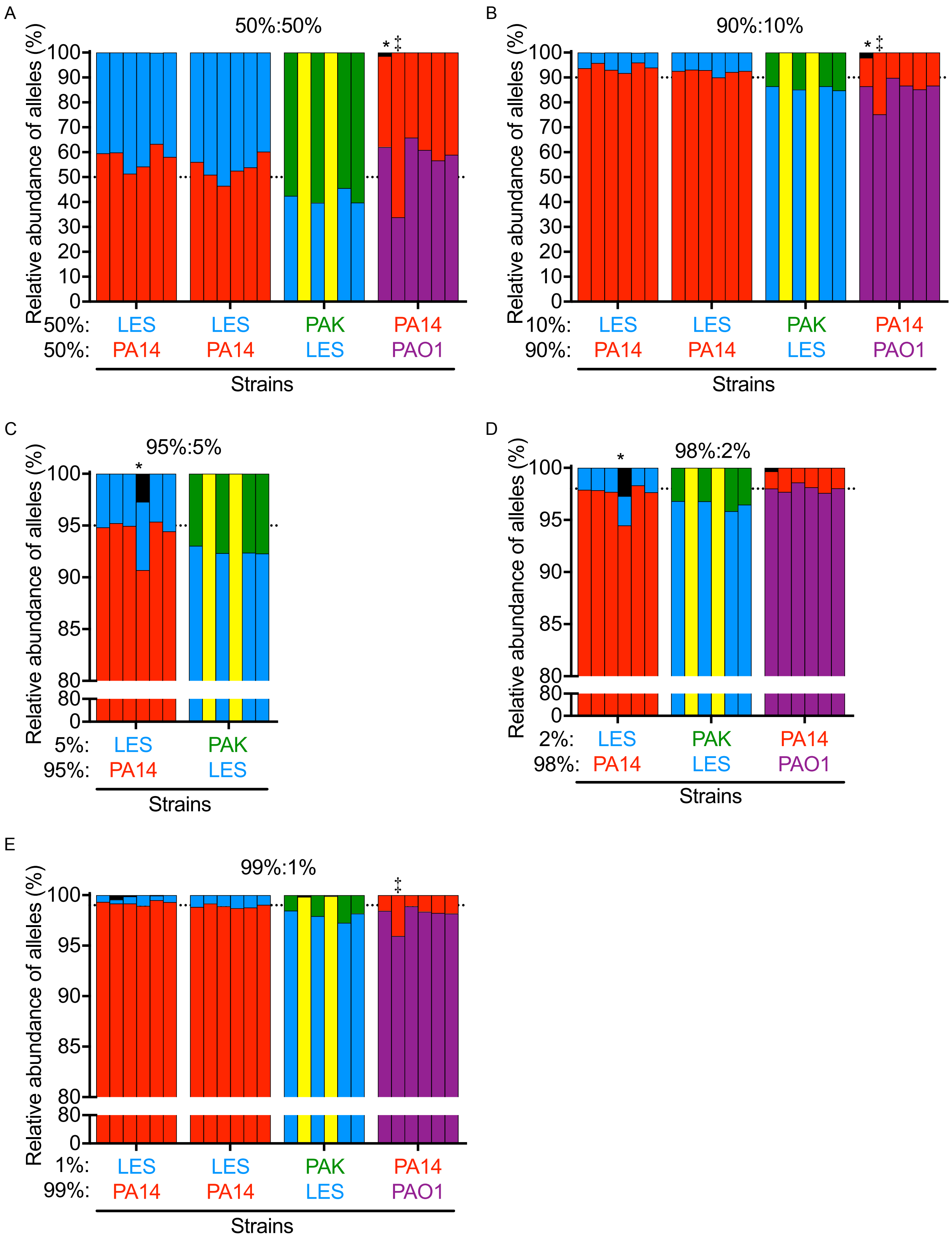
A
50%:50%
B
90%:10%
C
95%:5%
D
98%:2%
E
99%:1%
Relative abundance of alleles (%)
Strains
50%: LES LES PAK PA14
50%: PA14 PA14 LES PAO1
10%: LES LES PAK PA14
90%: PA14 PA14 LES PAO1
5%: LES PAK
95%: PA14 LES
2%: LES PAK PA14
98%: PA14 LES PAO1
1%: LES LES PAK PA14
99%: PA14 PA14 LES PAO1

**Figure S2. PopMLST on mixtures of Pa laboratory strains reproducibly identifies the relative abundance of each strain.** We tested pairwise combinations of four Pa strains with different MLST types across a range of ratios. In all cases, the two strains were identified in ratios close to expected. Each strain is represented by a different color (PAO1=purple, PA14=blue, LES=red, and PAK=green). PA14 and PAK have the same sequence for two loci indicated in yellow. PopMLST was performed on mixtures of cultured colonies for the PAO1:LES experiments, all others were performed on mixtures of DNA. Each set of bars represents an independent experiment. MLST alleles identified but not present in the mixtures, likely due to sequencing error, are indicated in black and those detected at >1%, are indicated with *. ‡ indicates PCR bias as evidenced by one allele being consistently under or overrepresented.

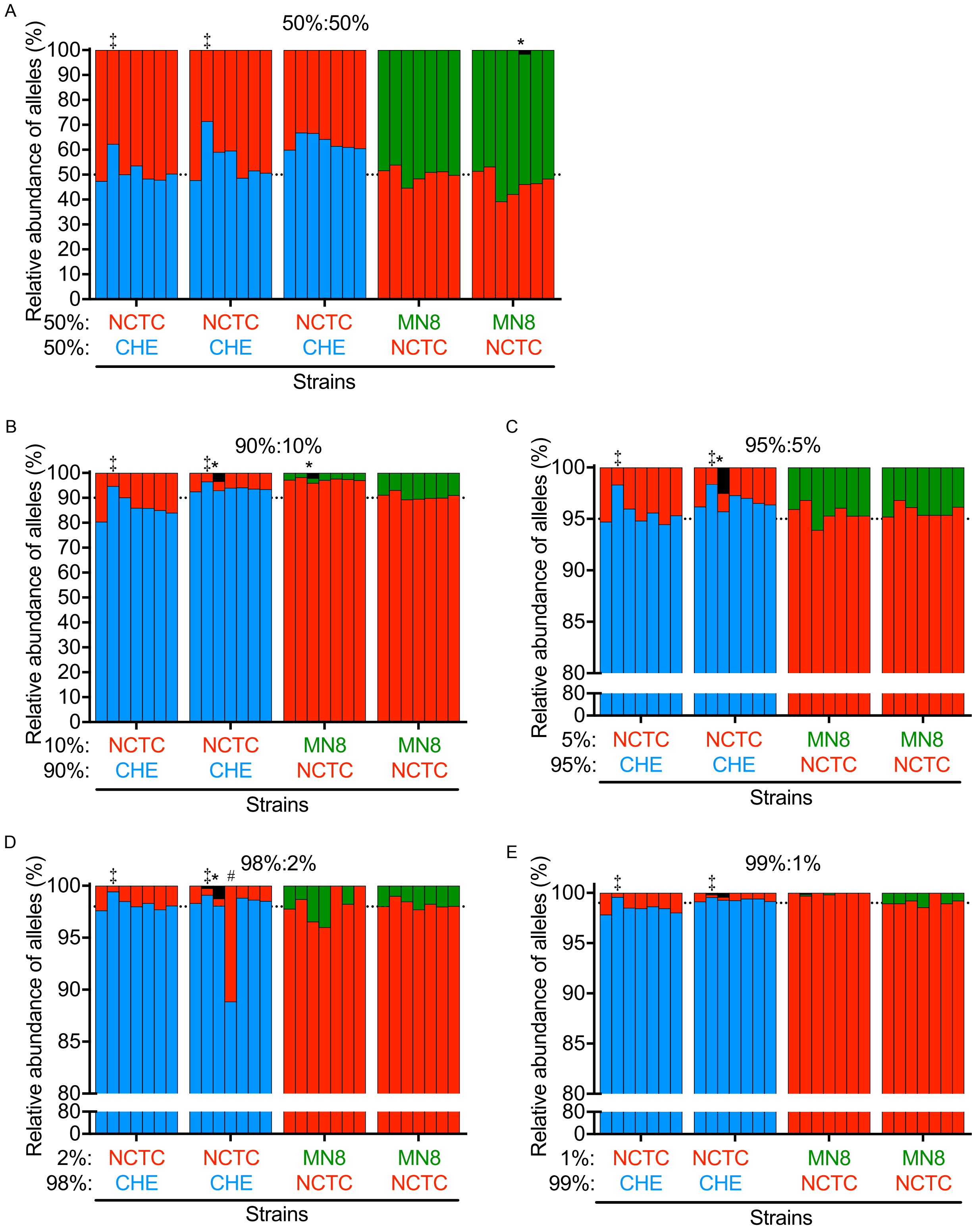


**Figure S3. PopMLST on mixtures of Sa laboratory strains reproducibly identifies the relative abundance of each strain.** We tested pairwise combinations of three Sa strains with different MLST types across a range of ratios. In all cases, the two strains were identified in ratios close to expected. Each strain is represented by a different color (NCTC8325=red, CHE=blue, MN8=green). Each set of bars represents an independent experiment. Black indicates the presence of a third allele. MLST alleles identified but not present in the mixtures, likely due to sequencing error, are indicated in black and those detected at >1%, are indicated with *. ‡ indicates PCR bias as evidenced by one allele being consistently under or overrepresented. # indicates non-systemic under/over representation of an allele, possibly due to jackpot amplifications.

Table S1. *S. aureus* strains and MLST types

| Strain | arc | aro | glp | gmk | pta | tpi | yqi | MLST type |
|---|---|---|---|---|---|---|---|---|
| NCTC8325 | 3 | 3 | 1 | 1 | 4 | 4 | 3 | 8 |
| CHE482 | 10 | 14 | 8 | 6 | 10 | 3 | 2 | 45 |
| MN8 | 2 | 2 | 2 | 2 | 6 | 3 | 2 | 30 |
| CF* | 13 | 13 | 1 | 1 | 12 | 11 | 13 | 15 |

* Clinical isolate from patient with Cystic Fibrosis

Table S2. *P. aeruginosa* strains and MLST types

| Strain | acs | aro | gua | mut | nuo | pps | trp | MLST type |
|---|---|---|---|---|---|---|---|---|
| PA14 | 4 | 4 | 16 | 12 | 1 | 6 | 3 | 253 |
| LES | 6 | 5 | 11 | 3 | 4 | 23 | 1 | 146 |
| PAO1 | 7 | 5 | 12 | 3 | 4 | 1 | 7 | 549 |
| PAK | 11 | 5 | 11 | 11 | 4 | 4 | 14 | 693 |
| CF5* | 28 | 5 | 30 | 3 | 3 | 4 | 14 | 1538 |
| CF6* | 40 | 5 | 11 | 5 | 4 | 38 | 37 | 167 |
| CF2* | 11 | 76 | 5 | 3 | 61 | 14 | 3 | 485 |

* Clinical isolate from patient with Cystic Fibrosis

Table S3. Primers used for PopMLST

| MLST loci | Published MLST primer | PopMLST primer |
|---|---|---|
| SaarcF | TTGATTCACCAGCGCGTATTGTC | TCGTCGGCAGCGTCAGATGTGTATAAGAGACAGCCGTTGATTCACCAGCGCGTATTGTC |
| SaarcR | AGGTATCTGCTTCAATCAGCG | GTCTCGTGGGCTCGGAGATGTGTATAAGAGACAGGCTAGGTATCTGCTTCAATCAGCG |
| SaaroF | ATCGGAAATCCTATTTCACATTC | TCGTCGGCAGCGTCAGATGTGTATAAGAGACAGCCGATCGGAAATCCTATTTCACATTC |
| SaaroR | GGTGTTGTATTAATAACGATATC | GTCTCGTGGGCTCGGAGATGTGTATAAGAGACAGGCTGGTGTTGTATTAATAACGATATC |
| SaglpF | CTAGGAACTGCAATCTTAATCC | TCGTCGGCAGCGTCAGATGTGTATAAGAGACAGCCGCTAGGAACTGCAATCTTAATCC |
| SaglpR | TGGTAAAATCGCATGTCCAATTC | GTCTCGTGGGCTCGGAGATGTGTATAAGAGACAGGCTTGGTAAAATCGCATGTCCAATTC |
| SagmkF | ATCGTTTTATCGGGACCATC | TCGTCGGCAGCGTCAGATGTGTATAAGAGACAGCCGATCGTTTTATCGGGACCATC |
| SagmkR | TCATTAACTACAACGTAATCGTA | GTCTCGTGGGCTCGGAGATGTGTATAAGAGACAGGCTTCATTAACTACAACGTAATCGTA |
| SaptaF | GTTAAAATCGTATTACCTGAAGG | TCGTCGGCAGCGTCAGATGTGTATAAGAGACAGCCGGTTAAAATCGTATTACCTGAAGG |
| SaptaR | GACCCTTTTGTTGAAAAGCTTAA | GTCTCGTGGGCTCGGAGATGTGTATAAGAGACAGGCTGACCCTTTTGTTGAAAAGCTTAA |
| SatpiF | TCGTTCATTCTGAACGTCGTGAA | TCGTCGGCAGCGTCAGATGTGTATAAGAGACAGCCGTCGTTCATTCTGAACGTCGTGAA |
| SatpiR | TTTGCACCTTCTAACAATTGTAC | GTCTCGTGGGCTCGGAGATGTGTATAAGAGACAGGCTTTTGCACCTTCTAACAATTGTAC |
| SayqiF | CAGCATACAGGACACCTATTGGC | TCGTCGGCAGCGTCAGATGTGTATAAGAGACAGCCGCAGCATACAGGACACCTATTGGC |
| SayqiR | CGTTGAGGAATCGATACTGGAAC | GTCTCGTGGGCTCGGAGATGTGTATAAGAGACAGGCTCGTTGAGGAATCGATACTGGAAC |
| PaacsF | GCCACACCTACATCGTCTAT | TCGTCGGCAGCGTCAGATGTGTATAAGAGACAGCCGGCCACACCTACATCGTCTAT |
| PaacsR | AGGTTGCCGAGGTTGTCCAC | GTCTCGTGGGCTCGGAGATGTGTATAAGAGACAGAGGTTGCCGAGGTTGTCCAC |
| PaaroF | ATGTCACCGTGCCGTTCAAG | TCGTCGGCAGCGTCAGATGTGTATAAGAGACAGCCAATGTCACCGTGCCGTTCAAG |
| PaaroR | GTTCCTTGGCTGACGGAAGT | GTCTCGTGGGCTCGGAGATGTGTATAAGAGACAGGGTTGAAGGCAGTCGGTTCCTTG |
| PaguaF | AGGTCGGTTCCTCCAAGGTC | TCGTCGGCAGCGTCAGATGTGTATAAGAGACAGCCAGGTCGGTTCCTCCAAGGTC |
| PaguaR | GACGTTGTGGTGCGACTTGA | GTCTCGTGGGCTCGGAGATGTGTATAAGAGACAGCGACGTTGTGGTGCGACTTGA |
| PamutF | AGAAGACCGAGTTCGACCAT | TCGTCGGCAGCGTCAGATGTGTATAAGAGACAGCCGAGAAGACCGAGTTCGACCAT |
| PamutR | GGTGCCATAGAGGAAGTCAT | GTCTCGTGGGCTCGGAGATGTGTATAAGAGACAGGCAGGGTGCCATAGAGGAAGTCAT |
| PanuoF | ACGGCGAGAACGAGGACTAC | TCGTCGGCAGCGTCAGATGTGTATAAGAGACAGCCACGGCGAGAACGAGGACTAC |
| PanuoR | TGGCGGTCGGTGAAGGTGAA | GTCTCGTGGGCTCGGAGATGTGTATAAGAGACAGTGGCGGTCGGTGAAGGTGAA |
| PappsF | GGTGACGACGGCAAGCTGTA | TCGTCGGCAGCGTCAGATGTGTATAAGAGACAGGGTGACGACGGCAAGCTGTAC |
| PappsR | GTATCGCCTTCGGCACAGGA | GTCTCGTGGGCTCGGAGATGTGTATAAGAGACAGGGTATCGCCTTCGGCACAGGA |
| PatrpF | TTCAACTTCGGCGACTTCCA | TCGTCGGCAGCGTCAGATGTGTATAAGAGACAGCTTTTCAACTTCGGCGACTTCCATGT |
| PatrpR | GGTGTCCATGTTGCCGTTCC | GTCTCGTGGGCTCGGAGATGTGTATAAGAGACAGCGGTGTCCATGTTGCCGTTCC |

Table S4. PCR conditions

| Loci | PCR reagent | Ta |
|---|---|---|
| *S. aureus* | | |
| arc | Q5 (NEB) | 61 |
| aro | Q5 (NEB) | 56 |
| glp | Q5 (NEB) | 56 |
| gmk | Phusion (NEB) | 56 |
| pta | Q5 (NEB) | 61 |
| tpi | Q5 (NEB) | 61 |
| yqi | Q5 (NEB) | 61 |
| *P. aeruginosa* | | |
| arc | Kapa (Roche) | 59 |
| gua | Kapa (Roche) | 59 |
| mut | Q5 + GC buffer (NEB) | 55 |
| nuo | Kapa (Roche) | 62 |
| pps | Q5 + GC buffer (NEB) | 67 |
| trp | Q5 + GC buffer (NEB) | 67 |
